# Design principles for selective polarization of PAR proteins by cortical flows

**DOI:** 10.1101/2022.09.05.506621

**Authors:** Rukshala Illukkumbura, Nisha Hirani, Joana Borrego-Pinto, Tom Bland, KangBo Ng, Lars Hubatsch, Jessica McQuade, Robert G. Endres, Nathan W. Goehring

**Affiliations:** The Francis Crick Institute, 1 Midland Road, London, UK; Institute for the Physics of Living Systems, University College London, London, UK; Department of Life Sciences, Imperial College London, London, UK

**Keywords:** Cell polarity, PAR proteins, actomyosin, cortical flow, advection, symmetry-breaking

## Abstract

Clustering of membrane-associated molecules is thought to promote interactions with the actomyosin cortex, enabling size-dependent transport by actin flows. Consistent with this model, in the *C. elegans* zygote, anterior segregation of the polarity protein PAR-3 requires oligomerization. However, through direct assessment of advection of PAR proteins, we not only find no links between PAR-3 advection and oligomer size, but also observe efficient advection of both anterior and posterior PAR proteins. Consequently, differential cortex engagement cannot account for selective size-dependent PAR protein transport. Instead, combining experiment and theory we demonstrate that segregation efficiency of PAR proteins by cortical flow is determined by the stability of membrane association, which is enhanced by clustering and specifies persistence of transport. Indeed, stabilizing membrane association was sufficient to invert polarity of a normally posterior PAR protein. Our data therefore indicate that advection of membrane-associated proteins is more pervasive than anticipated and thus cells must tune membrane association dynamics to achieve differential transport by cortical flows.

## Introduction

Over half a century ago, it was observed that crosslinking of antigens on the surface of immune cells could induce their coalescence into a domain at the rear of immune cells (Raff et al., 1970). This process of “capping” results from the rearward transport of cross-linked antigens by retrograde cortical actin flows linked to cell motility (Bray and White, 1988; Taylor et al., 1971). However, in the years since there has been persistent debate over precisely what membrane-associated components are flowing together with the actin cortex, leaving unclear the principles that define whether a particular molecule will be subject to long range transport by cortical flows (Sheetz et al., 1989; Holifield et al., 1990; Jacobson and Kapustina, 2019; Bretscher, 1996; Illukkumbura et al., 2020).

Polarization of the *C. elegans* zygote initially proceeds similarly to the process of antigen capping. Here cortical actomyosin flows away from what will become the zygote posterior pole (Munro et al., 2004). In doing so, cortical flows segregate a conserved set of peripheral membrane-associated polarity proteins, PAR-3, PAR-6, and PKC-3 (aPKC) into a cap at what will become the anterior pole (Munro et al., 2004; Goehring et al., 2011 b). As these PAR proteins are segregated, they are replaced on the posterior plasma membrane by a second set of PAR proteins (PAR-1, PAR-2, LGL-1, and CHIN-1) (Cuenca et al., 2003; Goehring, 2014; Rose and Gonczy, 2014). Once flows cease, mutually antagonistic interactions between these anterior (aPAR) and posterior (pPAR) proteins maintain the resulting polarity domains independently of the actin cortex (Cuenca et al., 2003; Goehring et al., 2011a; Goehring et al., 2011b; Hao et al., 2006; Hoege et al., 2010; Kumfer et al., 2010; Lang and Munro, 2017).

Similar to capping, segregation of aPAR proteins depends critically on plasma-membrane associated clusters of PAR-3. PAR-3 cluster motion is tightly coupled to the actin cortex and disruption of a conserved N-terminal oligomerization domain prevents polarization (Dickinson et al., 2017; Munro et al., 2004; Rodriguez et al., 2017; Wang et al., 2017). Because the membrane association of PAR-6 and PKC-3 is tightly linked to interactions with PAR-3, under normal conditions all three co-segregate (Beers and Kemphues, 2006; Rodriguez et al., 2017). However, when allowed to load onto the membrane independently of PAR-3, PAR-6 and PKC-3 fail to segregate (Rodriguez et al., 2017). Thus, the question arises: what is special about PAR-3 clusters that allows them to be so efficiently transported across length scales reaching tens of microns?

A key requirement for transport of molecules by cortical flow is that their motion be entrained by that of the underlying actomyosin cortex, a process referred to as advection. An attractive model for explaining cluster-dependent transport is that clustering results in an increase in effective friction arising from size-dependent binding to or corralling of clusters by the flowing actin cortex (Chang and Dickinson, 2022; Zmurchok and Holmes, 2022), similar to what has been proposed at the immunological synapse (Hartman et al., 2009). In such a model clustering would effectively tune the physical coupling of molecules to the actomyosin cortex, allowing molecules to selectively ‘sense flows.’ Consistent with such a model, PAR-3 diffusive mobility is restricted by the actin cortex in embryos (Sailer et al., 2015) and recent work has suggested a strict size threshold for directional transport by cortical flows (Chang and Dickinson, 2022). At the same time, clustering of molecules could also serve to reduce diffusivity and enhance membrane association via avidity effects, both of which would also be expected to enhance the efficiency of transport by cortical flow. Consequently, resolving how molecules can be selectively transported by cortical actomyosin flows requires that we directly assess the relative contributions of advection, diffusion, and membrane association, a task which is particularly challenging for molecules that are not obviously clustered or that may diffuse extensively at the membrane.

Here we develop a workflow to directly characterize the mobility of distinct pools of PAR proteins including clustered and non-clustered variants based on single particle tracking in live embryos. Surprisingly, our data are inconsistent with models of size- or cluster-dependent advection. Instead, we find that segregation of PAR proteins by cortical flows is largely defined by the strength of membrane binding, which increases the timescale over which cortical flows act. Consistent with this paradigm, increasing the strength of membrane binding of the normally posterior protein PAR-2 was sufficient to drive efficient anterior segregation and thus invert its polarity. Thus oligomerization-dependent tuning of membrane association represents a key design principle for achieving selective segregation by cortical flows in polarized cells.

## Results

### Advection of PAR-3 is independent of oligomeric state and diffusivity

To directly quantify the advection of individual molecules, we developed a single particle workflow that allowed us to measure particle displacements relative to the local actomyosin flow field (Figure 1A). Actomyosin generally flows from posterior to anterior. However, to account for local variations in the flow vector, we simultaneously image the molecule of interest along with cortical actomyosin to obtain the local flow vector (Figure 1B). Displacements of the molecule of interest are then projected onto local vectors parallel (x) and orthogonal (y) to flow. The resulting distributions are then fit to obtain an effective advection velocity along each axis, U_x_, U_y_, where U_y_ - the mean displacement orthogonal to flow - should be zero. The ratio of U_x_ to the velocity of the underlying actomyosin flow, v, defines a coupling coefficient, cc, where cc = U_x_/V. While this approach can in principle be used to analyze the motion of a single particle given a sufficient number of steps, in practice, imaging and tracking constraints, including photobleaching, mean that steps from a population of particles often must be pooled to obtain sufficient data. Validation of this workflow on known actomyosin cortex-associated molecules and simulated datasets confirmed our ability to reliably apply our method *in vivo* and suggests that we should be able to detect advection for effective diffusivities of < 0.3 μm^2^/s (Figure 1C-I), which is well in excess of the typical diffusivities that have been measured for PAR proteins (Arata et al., 2016; Goehring et al., 2011a; Gross et al., 2019; Hubatsch et al., 2019; Robin et al., 2014).

**Figure 1.**
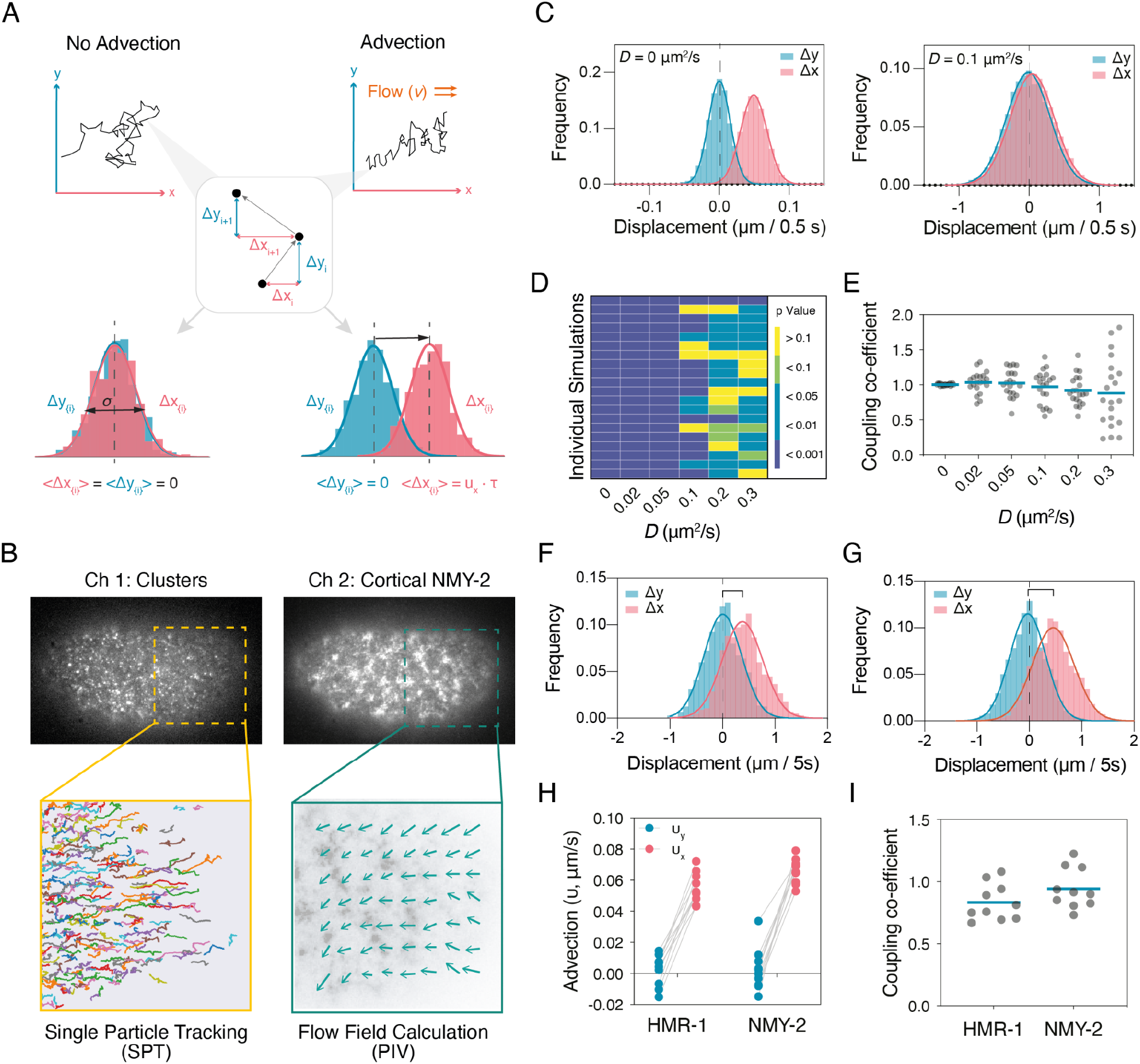
Extracting advection and diffusion parameters from single particle trajectories. **(A)** Schematic for decomposing advective and diffusive components to particle trajectories. Particle displacements are projected onto axes x and y defined as parallel and perpendicular to the local flow vector, respectively. In principle these should take the form of a Gaussian. Drift is defined by the shift of the mean population displacement (<Δx>,<Δy>), along the relevant axis, where we expect <Δy> ~ 0. Advection velocity (U_x_) is then given by mean displacement <Δx>, divided by the time lag, τ, with the cortex coupling coefficient given by ratio of advection velocity (U_x_) to local flow velocity (CC = U_x_/V). **(B)** Schematic for extraction of particle motion (PAR-3 clusters shown) and local flow field. Two channel image series were captured for cortical NMY-2 and the molecule of interest. The resulting image series were subject to either a Python-based particle tracking scheme or particle image velocimetry (PIVLab, Matlab). Particle displacements for a given τ were then projected onto the relevant x- and y-axes defined by the local flow vector. Note, positive movement on the x-axis generally reflects motion *towards the anterior*. **(C)** Distribution of displacements in x and y for simulations of varying D for τ = 0.5 s. **(D)** Reliability of detection of drift as a function of *D*. Significance of difference between <Δx> and <Δy> calculated using 1000 random displacements (τ = 0.5 s). Results shown for 20 independent simulations. Student’s t-test, unpaired, two-tailed. **(E)** Mean CC of ~ 1.0 is obtained for all *D*, though error increases with *D*. Each point indicates CC measured from 1000 random displacements from a single simulated dataset in (D), with mean indicated. **(F-G)** Example of the distribution of displacements for (F) NMY-2 and (G) HMR-1 for single embryos (τ = 5s, n = 1000 randomly selected steps). In both cases, there is a characteristic drift component along the flow axis (x). Displacements parallel (red) and orthogonal (blue) to the local flow axis are shown. **(H)** Fit values for advection velocity for NMY-2 and HMR-1 for parallel (U_x_, red) and orthogonal (U_y_, blue) displacements relative to the flow axis. Lines connect paired data points from single embryos. **(I)** Both HMR-1 and NMY-2 exhibit coupling coefficients near 1. Mean values for individual embryos shown along with mean of all embryos.

To directly assess size-dependent effects on diffusion and advection of PAR-3 clusters, we tracked individual PAR-3 clusters during the polarity establishment phase. Taking mean cluster intensity as a proxy for size, we find as expected that diffusivity decreased with cluster size, though the changes were surprisingly small (Figure 2A, B). Overall, cluster displacements exhibited Gaussian distributions consistent with lateral diffusion in the membrane, but with a clear shift in the mean displacement parallel to the flow vector, x (Figure 2C, 2D). Strikingly, fitting revealed nearly identical coupling coefficients across all size bins that were only modestly reduced relative to the NMY-2 reference (Figure 2E). Thus, at least for the range of cluster sizes we observe and track, both the diffusion and advection of clusters is relatively insensitive to cluster size. We also re-binned particles by diffusivity to determine if slower particles might be advected better, reasoning that reduced diffusivity could reflect stronger coupling to or local corralling by the cortex. However, again we found that coupling coefficients were effectively unchanged for varying diffusivities (Figure 2F).

**Figure 2.**
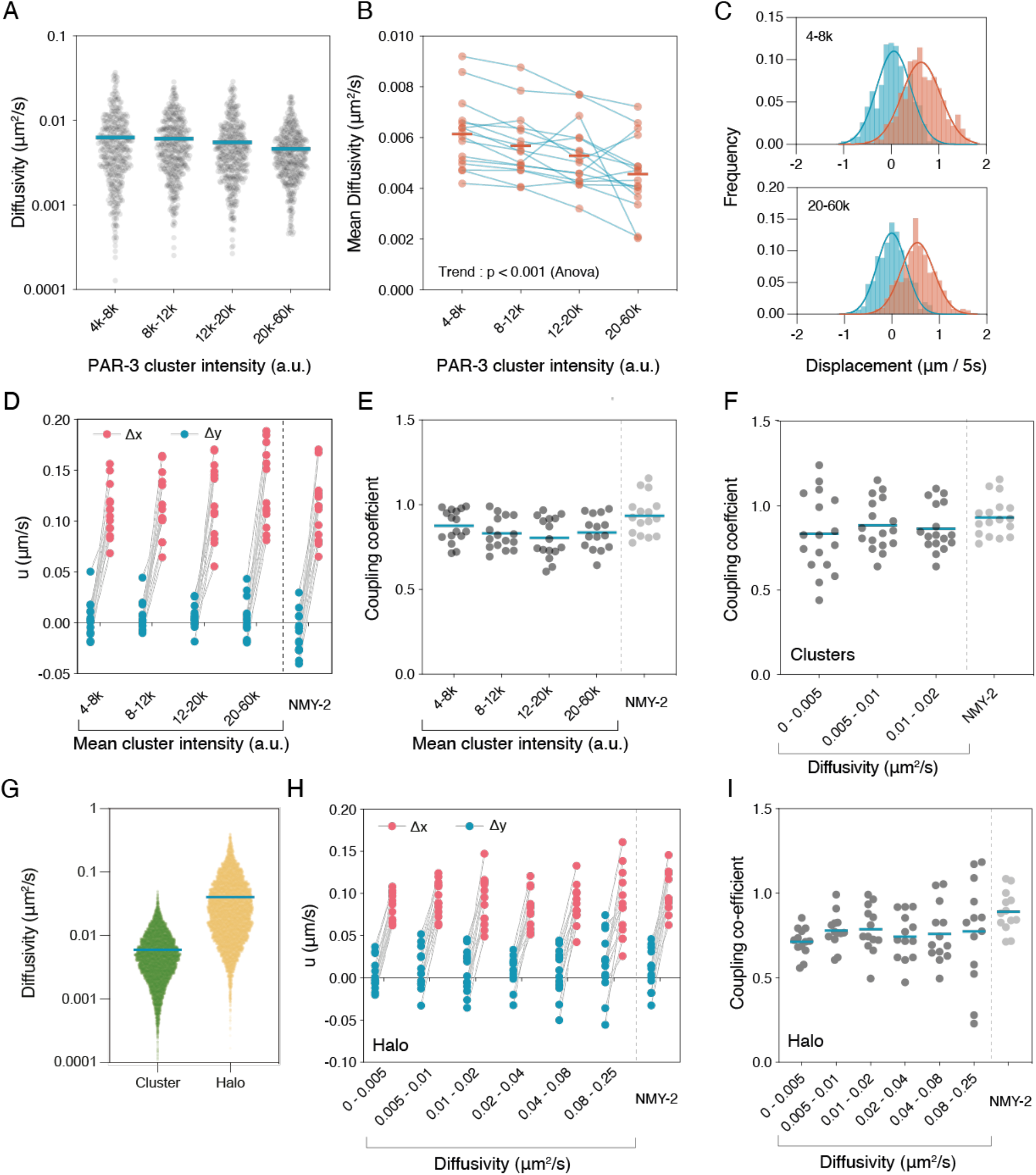
Advection of PAR-3 is independent of cluster size. **(A-B)** Cluster size only modestly affects diffusivity. (A) Distribution of diffusivities for 400 random clusters selected from all particles (16 embryos) for each of the indicated size bins. τ = 2 s. Mean values indicated. **(B)** Plot of mean cluster diffusivity per embryo across cluster size bins reveals a general trend of decreasing diffusivity with cluster size, but note that clusters are generally very slowly diffusing in all bins. Mean values across all embryos indicated by bars. **(C)** The distribution of PAR-3 cluster displacements are shifted along the flow-parallel axis. Data shown for all clusters from a single embryo, τ = 5s, n (steps) = 542, 1033, 1196, 844. **(D)** Fit values for advection velocity parallel (U_x_, red) and perpendicular (U_y_, blue) relative to the flow axis are similar for different sized clusters. Lines connect paired values from single embryos. **(E-F)** Coupling coefficients are independent of cluster size, diffusivity. Coupling coefficients were fit from particles binned by indicated size (E) or diffusivity (F) on a per embryo basis. Mean indicated. **(G)** Substoichiometric labeling of Halo::PAR-3 captures a faster diffusing PAR-3 pool. Distribution of effective diffusivities for PAR-3 clusters (green, n = 9996, 16 embryos) and single JF-549 labeled molecules (yellow, n = 12810, 12 embryos). Due to different imaging regimes, τ = 2 s (clusters), 0.2 s (Halo). Note log scale, mean indicated. **(H)** Measured parallel (U_x_, red) and perpendicular (U_y_, blue) advection velocities for single Halo-tagged molecules binned by diffusivity (t - 0.5 s). **(I)** Coupling coefficients derived from Halo::PAR-3 single molecule tracking are independent of diffusivity. Coupling coefficients were fit from molecules binned by indicated diffusivity on a per embryo basis. Mean indicated.

A caveat to these results is that imaging conditions optimized for clusters may fail to detect smaller, faster-diffusing species. Therefore, to better capture the full population and assess advection as a function of diffusivity, we turned to a single molecule labeling regime in which sub-stoichiometric labeling of endogenously Halo-tagged proteins was used to achieve a sparse set of well-separated trajectories that should better sample the overall population. The resulting distribution of diffusivities for Halo-tagged PAR-3 was both broader and shifted towards faster diffusivity compared to clusters (Figure 2G), indicating that we captured a substantial population of molecules with D > 0.02 um^2^/s, which were exceedingly rare in our cluster dataset. After binning particles by diffusivity, we again measured the distribution of displacements. Strikingly, advection of particles for even the fastest diffusing bins (D > 0.02 μm^2^/s), which reflect molecules missed by the cluster imaging regime, remained high and effectively unchanged compared to slower diffusing bins (2H, I). This was despite the fact that these molecules exhibited an effective diffusivity of up to an order of magnitude larger than is typical for clusters.

Finally, it remained possible that even the smallest particles we could detect at the membrane were oligomers of some form and that formation of a minimal oligomer is nonetheless required for advection, which would be consistent with (Chang and Dickinson, 2022). Oligomerization depends on the conserved CR1 oligomerization domain (Benton and Johnston, 2003; Feng et al., 2007; Mizuno et al., 2003; Zhang et al., 2013), mutation of which disrupts PAR-3 localization and leads to polarity defects and embryonic lethality (Dickinson et al., 2017; Li et al., 2010). We therefore introduced a small, previously characterized deletion into the endogenous par-3 locus to generate an oligomerization defective PAR-3, PAR-3(Δ69-82) (Figure 3A). Similar to prior results using this or similar mutations, introduction of the Δ69-82 mutation resulted in failure of PAR-3 to segregate into the anterior membrane and an overall loss of membrane association (Figure 3B). Consistent with defects in polarity, embryos exhibited high rates of symmetric P0 divisions, embryonic and adult lethality and sterility. Notably, par-3(Δ69-89) embryos exhibited partially reduced cortical flow, consistent with the fact that aPARs are required for actomyosin contractility (Figure 3B-F)(Dickinson et al., 2017; Li et al., 2010; Munro et al., 2004; Rodriguez et al., 2017).

**Figure 3.**
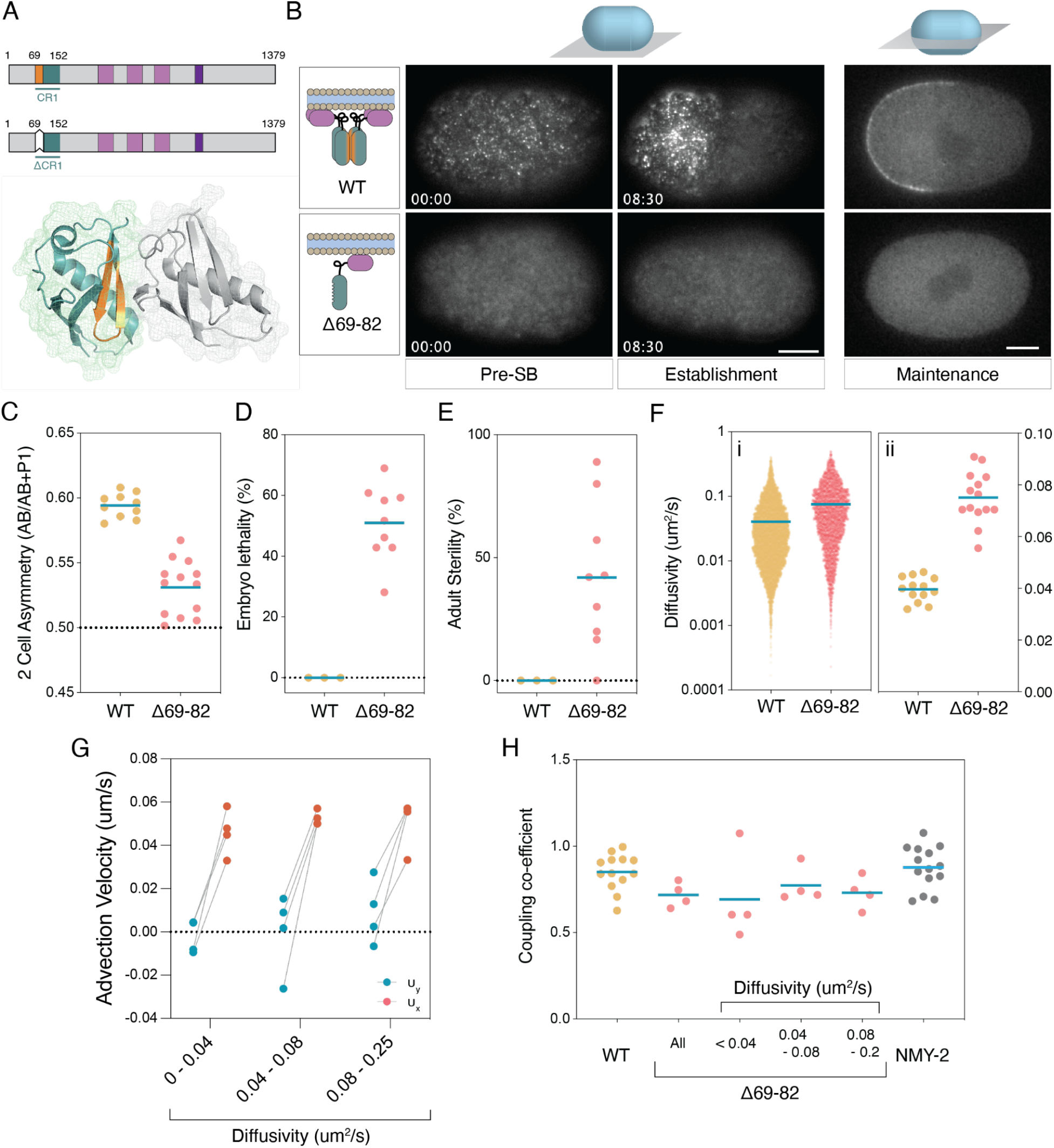
Advection of PAR-3 is independent of clustering. **(A)** Schematic of full length PAR-3 and PAR-3(Δ69-82) and a crystal structure of the PAR-3 CR1 domain dimer (rat PAR3-NTD, PDB:4I6P) are shown. Individual CR1 molecules are shown in grey, blue. Amino acids 69-82 (orange) are located at the oligomerization interface and color-coded in the schematics. **(B)** PAR-3(Δ69-82) does not accumulate on the anterior plasma membrane. Embryos expressing mNG fusions to either PAR-3(WT) or PAR-3(Δ69-82) are shown in a cortical plane by TIRF microscopy (Pre-Symmetry-Breaking, Establishment) or at midplane (Maintenance). Note general lack of signal at the plasma membrane, consistent with the requirement of oligomerization for stable membrane association. Scale bar = 10μm. Time mm::ss relative to symmetry breaking. **(C-E)** PAR-3(Δ69-82) leads to loss of P0 division asymmetry, embryonic lethality and sterility. (C) Daughter cell asymmetry (Area^AB^/Area^AB + P1^). (D) Percentage of embryos reaching L4 stage. (E) Surviving adults exhibiting sterility. Individual embryos (C) or replicates (D-E) shown with mean indicated. **(F)** PAR-3(Δ69-82) exhibits increased diffusivity relative to PAR-3(WT). HaloTag labeling was used to sparsely label PAR-3(Δ69-82) and effective diffusivity calculated from mean perpendicular step size for τ = 0.2 s (3616 particles, 14 embryos). Distribution of diffusivities for all molecules, mean indicated (i) and mean diffusivity per embryo, means indicated (ii). WT data reproduced from Figure 2G for comparison). **(G)** PAR-3(Δ69-82) displacements are biased along the flow axis. Plots of per-embryo mean displacements parallel and perpendicular to the flow axis for a representative embryo for τ = 0.5 s. **(H)** PAR-3(Δ69-82) couples to cortical flow. Coupling coefficients for PAR-3(Δ69-82) shown for all molecules or molecules binned by diffusivity shown compared to PAR-3(WT) and NMY-2. Datapoints for PAR-3(WT) and NMY-2 represent single embryos. Due to reduced numbers of molecules per embryo for PAR-3(Δ69-82) embryos, each data point in PAR-3(Δ69-82) bins reflects molecules combined from multiple embryos to obtain sufficient numbers for fitting.

While membrane association was reduced to the point of being nearly undetectable over cytoplasmic background when using a mNG::PAR-3(Δ69-82) fusion, single molecule analysis using sparse labeling of a Halo-tagged allele allowed us to track sufficient numbers of membrane binding events for analysis (Movie S1). Overall PAR-3(Δ69-82) molecules exhibited faster mean diffusivity compared to wild-type PAR-3, consistent with reduced clustering (Figure 3G). Strikingly, we still detected a clear signature of advection (Figure 3H). While slightly reduced compared to wild-type, coupling coefficients remained high and consistent over all diffusive bins analyzed (Figure 3I). Thus, the failure of PAR-3(Δ69-82) to exhibit significant segregation is not due to a failure to be advected and indicates that the ability of PAR-3 to couple to cortical flows over short timescales is unaffected by its oligomeric state or effective diffusivity.

### Rescue of cluster-defective PAR-3

We next asked whether we could rescue the segregation of monomeric PAR-3(Δ69-82) through replacing CR1-dependent oligomerization either with dimeric (2mer) or tetrameric (4mer) forms of the GCN4 leucine zipper to restore clustering or with an ectopic membrane targeting signal (RitC) to restore membrane localization independently of clustering (Figure 4A). All fusions exhibited normal advection (Figure 4D, Movie S2).

**Figure 4.**
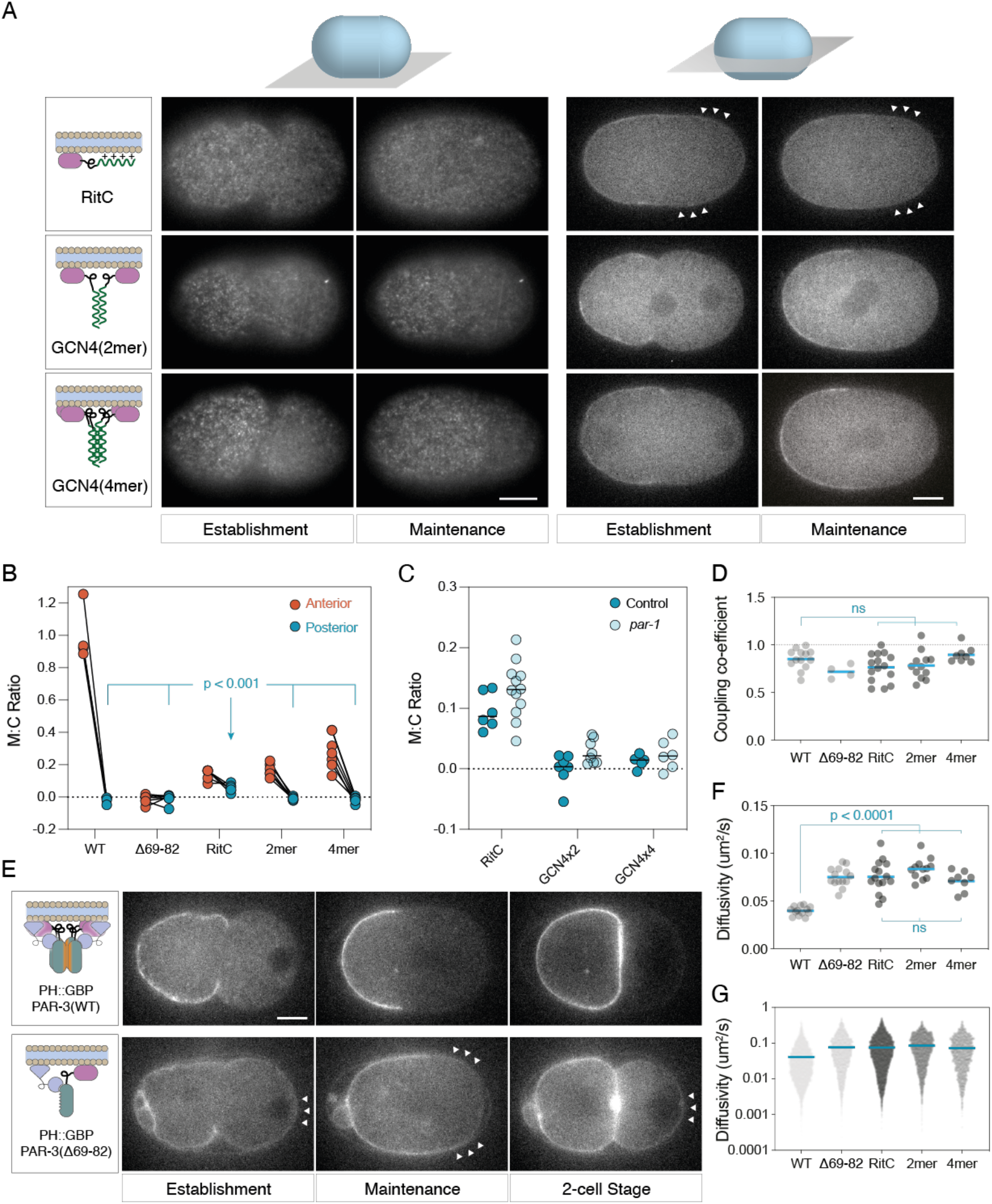
Rescue of monomeric PAR-3 segregation by ectopic membrane targeting and/or clustering. **(A)** (left) Schematic of rescue constructs fused to mNG::PAR-3(Δ69-82) shown along with example images of embryos taken at the cortex or at midplane at establishment and maintenance phases. Arrowheads indicate regions of clearance. Arrowheads indicate regions with posterior localization (RitC). Note midplane images are shown with identical scaling to highlight relative levels, while cortex images have been individually scaled to improve visibility of the clusters at the cortex. Scale bars = 10 μm. **(B)** Quantification of membrane to cytoplasm (M:C) ratio at the anterior and posterior pole at maintenance phase highlights reduced ability of PAR-3(RitC) to remain excluded from the posterior pole compared to clustered variants (Tukey’s multiple comparison test). **(C)** Posterior clearance is unaffected by *par-1(RNAi*). M:C ratios at the posterior pole were calculated from embryos subject to *ctl* or *par-1(RNAi*). Mean values indicated. Differences between *ctl* and *par-1* were not significant, unpaired t-test, Holm-Šídák correction. **(D)** Coupling coefficients are similar for all variants. Coupling coefficients calculated for all molecules in individual embryos are shown with means. Tukey’s multiple comparison test - variants are not significantly different from WT or between themselves. **(E)** Effect of clustering on membrane tethered PAR-3. GFP fusions to either PAR-3(WT) or PAR-3(Δ69-82) were tethered to the membrane via a membrane associated nanobody (PH::GBP) and their ability to segregate in response to flows tested. Note that PAR-3(WT) segregates normally and is fully excluded from the posterior pole despite constitutive targeting to the plasma membrane and is absent from the posterior P1 daughter cell. By contrast, PAR-3(Δ69-82) is less efficiently cleared and partially reinvades the posterior once flows cease such that it is also present on the plasma membrane of the P1 daughter cell. Arrowheads highlight posterior GFP signal. Scale bar = 10 μm. **(F-G)** All PAR-3 variants exhibit similar increases in diffusivity relative to PAR-3(WT). HaloTag labeling was used to sparsely label PAR-3 variants and effective diffusivity calculated from mean perpendicular step size for τ = 0.2 s. Mean diffusivity per embryo (F) and for all molecules (G) shown with mean indicated. WT and PAR-3(Δ69-82) data reproduced from Figure 3F for comparison. Mean diffusivity per embryo values for variants are significantly different from wild type, but not from each other. Tukey’s multiple comparison test.

We found that simply stabilizing membrane binding via RitC was sufficient to allow some PAR-3 asymmetry to develop, though segregation was noticeably less efficient than PAR-3(WT) as levels on the posterior membrane remained significantly above zero (Figure 4B). Nonetheless, PAR-3(RitC) was able to rescue division size asymmetry, cortical flow velocity, and embryonic lethality observed in *par-3(Δ69-82*) embryos, though only partially rescued adult sterility (Figure S1).

By contrast, PAR-3(2mer) and PAR-3(4mer) exhibited highly efficient segregation, with effectively no detectable signal remaining in the posterior half of the embryo (Figure 4B). PAR-3(4mer) appeared noticeably more punctate and exhibited higher membrane concentrations compared to PAR-3(2mer) consistent with a larger oligomer size, though still substantially below that of PAR-3(WT)(Figure 4A). Division asymmetry, cortical flow rates, embryonic lethality, and adult sterility appeared normal in both 2mer and 4mer variants (Figure S1). Thus, ectopic dimerization of monomeric PAR-3 mutants is sufficient to rescue PAR-3 function.

To determine whether this difference in behavior between PAR-3(RitC) and oligomeric forms of PAR-3 (i.e. WT, 2mer, 4mer) could simply be due to differential sensitivity to posterior membrane exclusion by the posterior PAR kinase PAR-1 rather than differential segregation by flow, we repeated our analysis in embryos depleted for PAR-1 by RNAi. We found that membrane concentrations were slightly elevated in *par-1(RNAi*) in all cases, but the relative efficiency of segregation of the different variants was unchanged (Figure 4C).

Finally, to confirm that ectopic membrane targeting per se does not compromise PAR-3 segregation behavior, we induced ectopic membrane targeting of GFP fusions to wild-type and monomeric PAR-3 by co-expressing a membrane-associated anti-GFP nanobody (PH::GBP)(Rodriguez et al., 2017). Membrane-tethered monomeric PAR-3(Δ69-82) behaved similarly to PAR-3(RitC), exhibiting modest segregation. By contrast, membrane-tethered PAR-3(WT) was segregated as efficiently as wild type (Figure 4E).

On the basis of these experiments, we conclude that direct membrane targeting via RitC or PH::GBP can rescue membrane accumulation of PAR-3(Δ69-82) and enable a degree of segregation, consistent with all forms of PAR-3 undergoing advection. At the same time, the reduced segregation of PAR-3(RitC) compared to oligomeric variants despite similar advection efficiency allows us to eliminate differential advection as a mechanisms for cluster-specific transport of PAR-3. Differential segregation is also unlikely to be due to cluster-dependent changes in diffusivity as both GCN4 and RitC fusions exhibited similar diffusivity (Figure 4F-G). Consequently, oligomerization of PAR-3 must alter molecular behavior in other ways to ensure highly efficient transport out of the posterior of the polarizing zygote.

### Both anterior and posterior PAR proteins are advected with similar efficiency

So far, our data indicate that the specificity of cortical-flow dependent segregation achieved by clustering of PAR-3 is not due to its effects on diffusion or physical linkage to the actin cortex: all species of PAR-3 examined were advected by flows with similar efficiency and molecules with similar diffusivity could differ in their ability to stably segregate. However, while differential linkage to the cortex could not explain the dependence of PAR-3 segregation on its oligomeric state, it remained possible that PAR-3 itself is unique among PAR proteins in its ability to be advected by cortical flows. Consistent with this hypothesis, we previously showed that segregation of PAR-6 and PKC-3 was dependent on their ability to associate with PAR-3 clusters (Rodriguez et al., 2017). Similarly, posterior PAR proteins are generally not strongly segregated into the anterior, even under conditions in which they are constitutively membrane-associated during the period of cortical flow (Folkmann and Seydoux, 2019; Hao et al., 2006; Rodriguez et al., 2017).

To determine whether other PAR proteins are advected by cortical flows and thus whether failure to be advected could explain their reduced ability to segregate compared to PAR-3, we applied our analysis to sparsely labeled Halo-tagged fusions to PAR-1, PAR-2 and PAR-6 during the polarity establishment phase. In all three cases, molecules could clearly be seen moving towards the anterior during the period of cortical flow (Figure 5A, Movie S3). Plots of displacements revealed a clear drift parallel to the local flow vector, consistent with all three proteins being advected by cortical flow (Figure 5B, 5C). For all three proteins, coupling coefficients were similar and generally exceeded 0.7 (Figure 5D). Both PAR-1 and PAR-2 exhibited higher mean diffusivity compared to PAR-6, which was itself significantly higher than that observed for PAR-3, indicating a general lack of correlation between diffusivity and the ability of molecules to couple to cortical flows (Figure 5E). Thus, despite their distinct behaviors in the embryo, an ability to link to and be advected by cortical flows is a general property of PAR proteins. Consequently, selective advection cannot account for the preferential ability of PAR-3 to be segregated by cortical flows.

**Figure 5.**
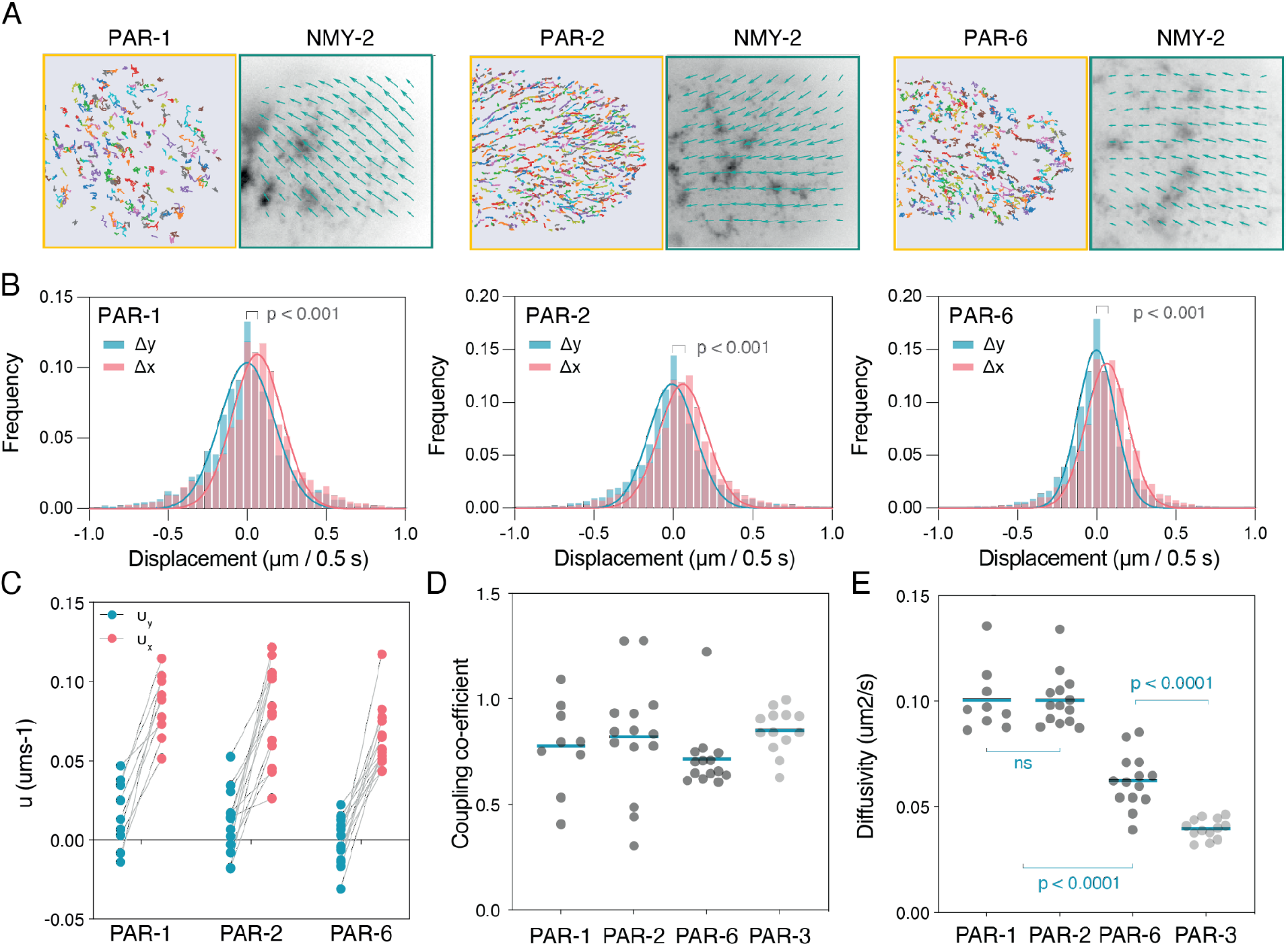
Anterior and posterior PAR proteins are advected with similar efficiency. **(A)** Sample trajectories (>15 frames) from a region in the embryo posterior shown for Halotag-labeled PAR-1, PAR-2, PAR-6. Corresponding flow fields derived from PIV analysis of NMY-2 shown at right. Vectors were normalized to highlight direction of vectors. **(B)** Plots of displacements parallel and perpendicular to the flow axis fit with a Gaussian distribution. **(C)** Fit values for advection velocity for PAR-1, PAR-2, and PAR-6 from displacements parallel (U_x_, red) and perpendicular (U_y_, blue) to the local flow axis highlight drift along the flow axis. Lines connect paired data from single embryos. **(D)** Coupling coefficients for PAR-1, PAR-2, and PAR-6 are similar to those observed for PAR-3. Individual embryo data points and mean shown. **(E)** Mean diffusivity per embryo for PAR-1, PAR-2, PAR-3, and PAR-6 highlight lack of correlation between diffusivity and advection (coupling coefficient). Note PAR-3 data reproduced from Figure 3F for comparison.

### A simple model for advective transport predicts segregation efficiency

To gain insight into how proteins may be efficiently segregated, we considered a simple model in which molecules undergo diffusion, membrane association/dissociation from a uniform cytoplasmic pool, and advection. Given the lack of effect of oligomer size on advection, the simplest explanation to explain differential segregation would be through effects of oligomerization on the diffusivity and/or membrane dissociation rate (k_off_). Generally, oligomerization should result in both a larger molecule, hence reduced diffusivity, and reduced membrane dissociation due to increased membrane avidity, both of which would generally be expected to enhance segregation.

Taking account of our results so far, we let molecules be advected equally and varied D and the membrane dissociation rate (k_off_) across multiple orders of magnitude. We did not impose any spatial regulation of these processes besides the imposition of a spatially varying cortical flow velocity that was fit to experimentally determined flow fields. We specifically did not include any feedback which could amplify or destabilize asymmetries. We applied an advection regime in which molecules were subject to advection for a period of 500 s, corresponding to the 5-10 minute polarity establishment phase during which flows initially pattern the PAR proteins (Blanchoud et al., 2015; Goehring et al., 2011b; Gross et al., 2019). Subsequently, to predict the persistence of asymmetry during the period after flows cease and before cytokinesis during which polarity is maintained, we then set advection to zero and allowed the system to equilibrate for a further 300 s. We scored both segregation efficiency, quantified by the peak magnitude of asymmetry (ASI - ASymmetric Index) and the relative posterior depletion after a 500 s period of cortical flow, as well as the persistence of asymmetry as measured by the timescale of asymmetry decay during the flow-independent maintenance phase.

As expected, segregation efficiency, measured either by ASI or posterior depletion, was maximized as k_off_ and D were reduced (Figure 6A, 6B), as was the stability of the induced asymmetry (Figure 6C, 6D). PAR proteins, however, typically exhibit diffusivities characterized by D <= ~ 0.1 μm^2^/s. In this regime, further reductions in D make relatively little difference and consequently, cluster-induced reductions in diffusivity of PAR-3 (e.g. from ~0.08 to <0.005 um^2^/s) would be expected to have only very minor effects on segregation efficiency (< 2%, Figure 6E-H). This result is consistent with diffusion being a poor predictor of differential segregation of our PAR-3 variants (see Figure 4). By contrast, in this regime, segregation is highly sensitive to k_off_. Notably, segregation is effectively non-existent for k_off_ > 0.1 s^-1^, while for k_off_ < 0.001 s^-1^, segregation is highly efficient and the resulting asymmetry decays only very slowly once flows cease (Figure 6E-G, 6I). Thus, in the absence of additional feedback and for typical diffusivities of PAR proteins, both the magnitude of asymmetry in response to cortical flows and its persistence once flows cease is predicted to be primarily set by how long molecules remain associated with the membrane.

**Figure 6.**
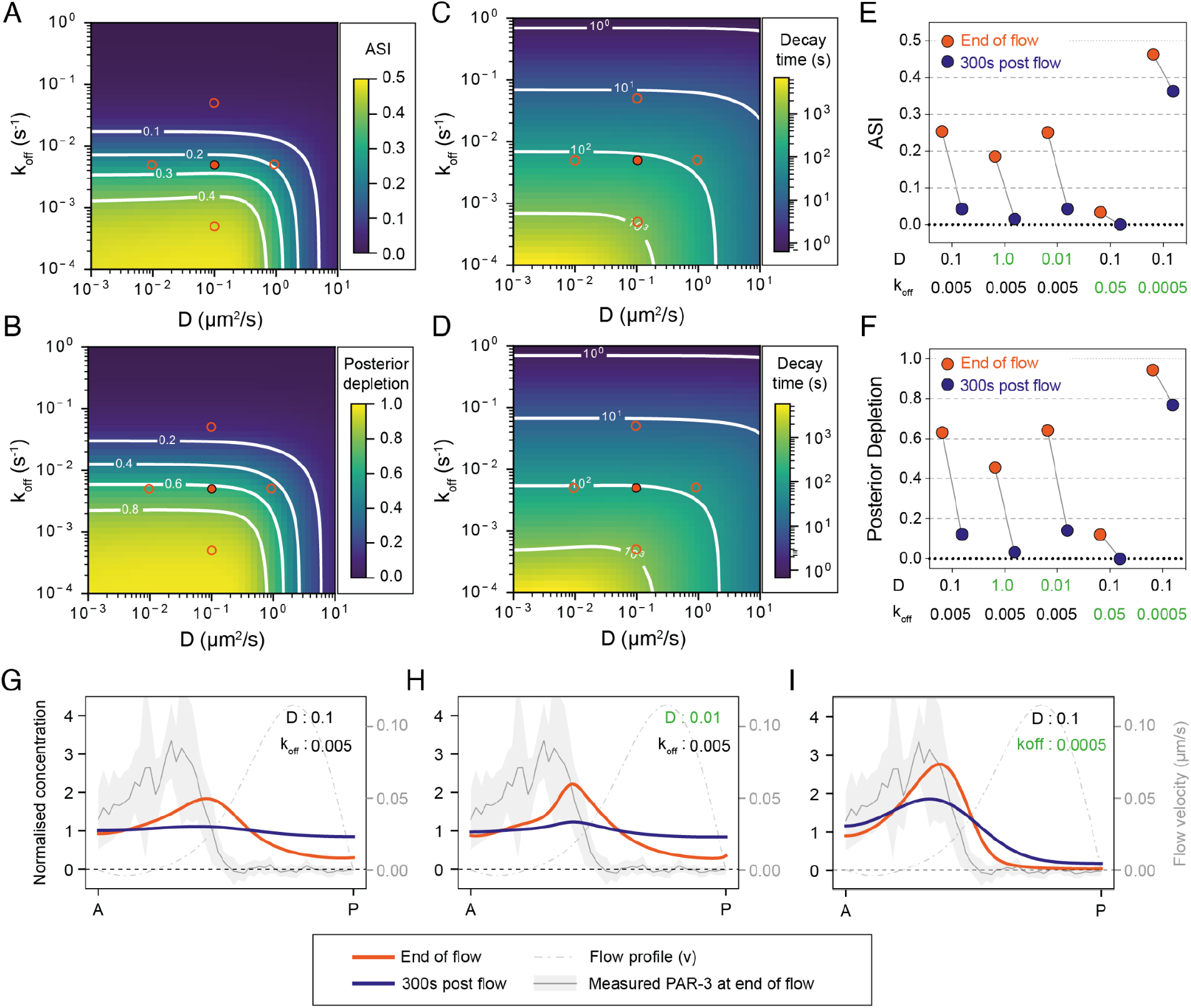
Predicted efficiency of advective segregation as a function of D and k_off_. **(A-D)** The efficiency of advective segregation of a hypothetical molecule as a function of k_off_ and D when subjected to an experimentally fit flow profile for a period of 500 s. (A-B) show maximum ASI and relative posterior depletion at the end of 500 s of flow, respectively. (C-D) show the characteristic time (t1/2) for decay in ASI and posterior depletion after flow ceases. Red circles in (A-D) indicate example parameters sets approximating the measured behavior of PAR-2 (solid circle, D ~ 0.1 μm^2^/s, k_off_ ~ 0.005 s^-1^) along with 10-fold increases or decreases in D and k_off_ (open circles). **(E-F)** Peak ASI (E) and posterior depletion (F) shown at the end of flow and 300 s later, equivalent to peak establishment phase and post NEBD maintenance phase, respectively for the example parameter sets denoted in (A-D). Note the magnitude and stability of asymmetry are strongly impacted by 10-fold changes in k_off_, but not similar changes in D. **(G-I)** Sample concentration profiles at the end of flow and 300s post flow for the indicated parameter sets from (A-F). The flow velocity function and measured PAR-3 distribution (mean ± SD) at the end of flow are shown for reference.

For the known membrane dynamics of PAR-2 and PAR-6 (D ~ 0.1 um^2^/s, k_off_ ~ 0.005 s^-1^) the model predicts modest and transient of segregation (ASI ~ 0.25, posterior depletion ~ 0.6, t1/2 ~ 100 s) (Figure 6A-G) (parameters from this work and (Goehring et al., 2011a; Hubatsch et al., 2019; Robin et al., 2014)). Thus, measurements of the segregation of PAR-2 and PAR-6 *in vivo* represent a key test of the model.

Because PAR-6 associates with PAR-3 clusters and PAR-3 is normally required for both PAR-6 membrane association and cortical flows, we could not measure PAR-3-independent segregation of PAR-6 in a wild-type context. However, association of PAR-6 and PKC-3 with PAR-3 clusters can be suppressed by inhibition of PKC-3 (Rodriguez et al., 2017). We find that in these conditions PAR-6 is still able to acquire some asymmetry, though to a substantially lower degree compared to when it associates with PAR-3 clusters. Further, unlike PAR-3, the resulting asymmetry decayed rapidly once flows ceased, consistent with predictions (Figure S2).

Similar to PAR-6, we cannot analyze PAR-2 segregation in wild-type conditions as it is actively excluded from the anterior by PKC-3. Thus, we introduced an alanine mutation into the key PKC-3 phosphosite, S241, which has been shown to prevent PKC-3-dependent displacement of PAR-2 from the plasma membrane and was previously reported to lead to uniform PAR-2 localization to the membrane (Hao et al., 2006; Motegi et al., 2011). We found that PAR-2(S241A) was in fact transiently enriched in the anterior by cortical flows, reaching a peak ASI of ~ 0.26 and peak depletion of PAR-2(S241A) in the posterior of ~0.56, figures that quantitatively matched the predictions of our model (Figure 7A - Establishment, B-D, Peak Flow). This asymmetry then dissipated substantially after flows ceased. Both ASI and peak depletion decayed by >70% within ~300s, again consistent with predictions of our model (Figure 7A - maintenance, B). Thus, the segregation behavior of PAR-2 and PAR-6 is fully consistent with predictions of this simple model of advective segregation.

**Figure 7.**
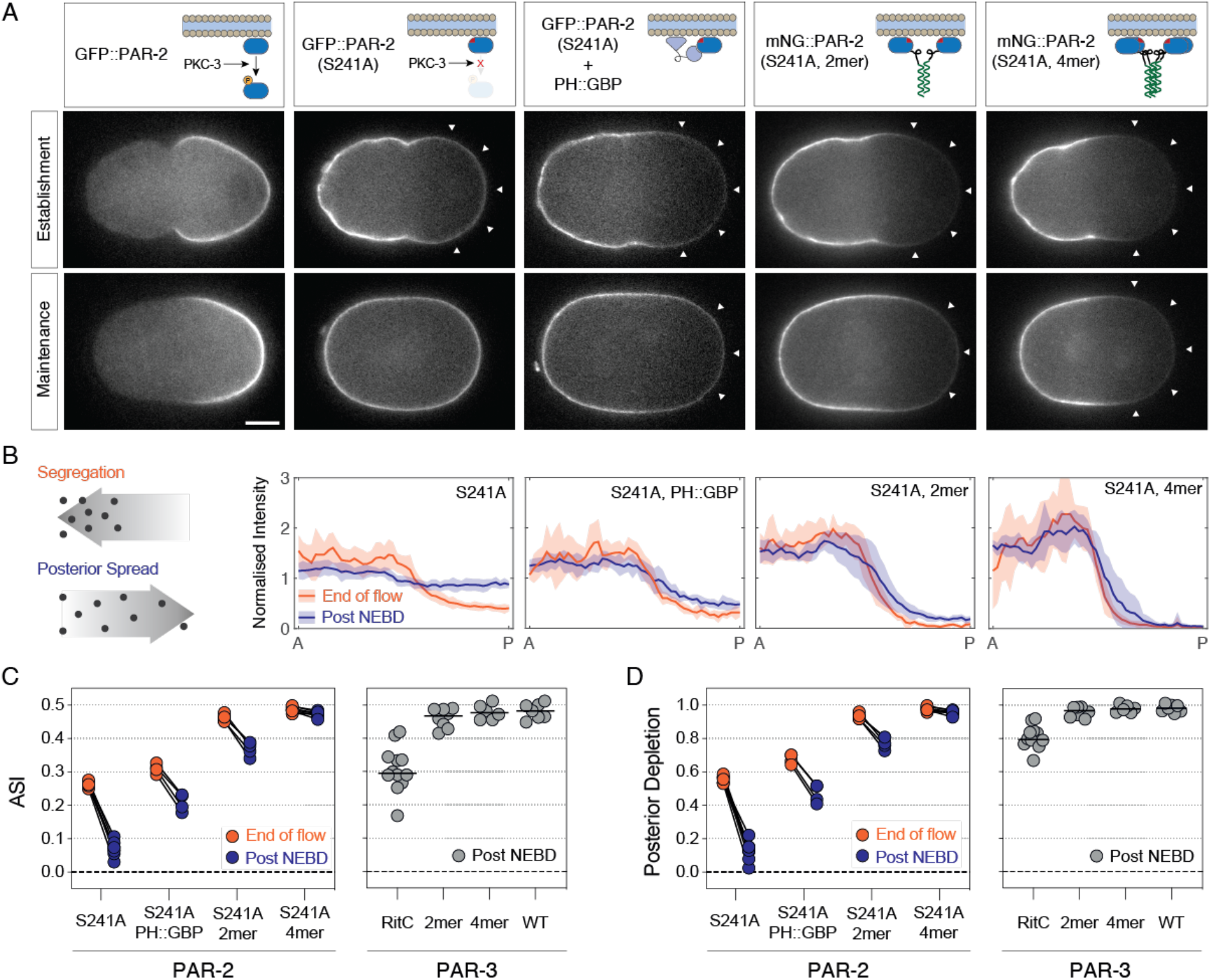
Increasing membrane avidity is sufficient to anteriorize posterior protein PAR-2. **(A)** Mutation of the PKC-3 target site in PAR-2, PAR-2(S241A) leads to modest transient anterior enrichment of PAR-2. Note reversed asymmetry compared to PAR-2(WT). The magnitude and stability of anterior segregation is enhanced by tethering to the membrane (PH::GFP) or by oligomerization (2mer, 4mer). Representative midplane confocal images captured at the end of the establishment phase (End of flow) and maintenance phase (Post NEBD) highlighting peak segregation and degree of asymmetry decay once flows cease. **(B)** Quantitation of membrane profiles of zygotes expressing PAR-2(S241A) variants shown in (A). Mean ± SD shown. **(C-D)** Quantification of asymmetry index (ASI) and relative posterior depletion for PAR-2(S241A) variants plotted along with Post NEBD data for PAR-3 zygotes for comparison (see Figure 4). Note that both oligomeric forms of PAR-2(S241A, 2mer/4mer) achieve near maximal levels for both ASI (0.5) and posterior depletion (1.0) during the establishment phase, which decay only minimally once flows cease. Data for S241A is pooled from GFP:: and mNG::PAR-2(S241A), which behaved identically.

### Tuning membrane binding is sufficient to ‘anteriorize’ posterior PAR protein PAR-2

If stability of membrane binding is the key determinant of efficient PAR segregation, one should be able to tune segregation efficiency by varying membrane affinity. To explicitly test this prediction, we stabilized membrane association of PAR-2(S241A) either by tethering it to the membrane via co-expression of PH::GBP with a GFP::PAR-2(S241A) fusion or by oligomerization, which would effectively increase the number of effective membrane binding domains as a function of oligomer size. In this case, we utilized the same dimeric (2mer) and tetrameric (4mer) forms of the GCN4 leucine zipper as we used in our PAR-3 experiments.

These modifications exhibited a progressive increase in both peak asymmetry of PAR-2(S241A) as measured by ASI and posterior depletion at the end of the polarity establishment phase, and in the persistence of this asymmetry once flows ceased in the polarity maintenance phase (Figure 7). Membrane targeting had the most modest effect, consistent with the lower membrane affinity of the PH_PLCΔ1_ domain relative to PAR-2 (k_off_ ~ 0.12 vs 0.0056 s^-1^)(Goehring et al., 2010; Goehring et al., 2011a). By contrast, both 2mer and 4mer forms of PAR-2 exhibited almost a complete reversal of PAR-2 polarity, attaining peak asymmetries of >0.46 (max = 0.5) and peak depletion in the posterior of >0.93 (max = 1.0). These values were comparable to that observed for oligomeric forms of PAR-3 (ASI > 0.47, Posterior Depletion > 0.97, see Figure 7D for comparison). Moreover, reversal of polarity was persistent, with ASI and posterior depletion decaying by less than 20% in the case of PAR-2(S241A, 2mer) and less than 5% in the case of PAR-2(S241A, 2mer) over the 5-6 minutes following the cessation of flows (Figure 7B-C). Indeed, the profile of tetrameric PAR-2 at the end of flows is conspicuously similar to PAR-3 (Compare Figure 7B, 241A, 4mer and Figure 6I). Thus, modulation of membrane binding appears sufficient to achieve efficient, persistent, and selective anterior segregation of PAR-2 by cortical flows, effectively reprogramming PAR-2 to achieve the pattern of polarization exhibited by oligomeric PAR-3.

## Discussion

While cortical flow-associated polarized transport has been observed for over a half century, the design features of molecules that determine whether molecules “go with the flow” have remained contentious. Experiments in non-adherent cells have proposed that cortical flows can induce bulk flow of plasma membrane components, including lipids, towards the cell rear leading to a treadmilling-like behavior. In such models, asymmetries arise from differential diffusion and/or membrane internalization (Bretscher, 1996; O’Neill et al., 2018; Tanaka et al., 2017). However, in other systems, advection appears selective with no evidence for lipid flow (Lee et al., 1990; Sheetz et al., 1989). In both systems, clustering could be expected to tune segregation. In the former, the primary effects of clustering would be envisioned to impact diffusivity or membrane turnover, while in the latter, clustering is thought to promote advection by altering the effective friction with the underlying cortex, for example through size-dependent entanglement with the cortical actin meshwork or other molecules linked to it.

Here we have shown that PAR proteins are indeed subject to differential transport by cortical flows in a manner that correlates with clustering. However, differential transport does not arise through changes in advective coupling to the actin cortex. Most notably, there is no obvious correlation between the advection of a molecule and either its oligomer size or diffusivity which one might expect would be reflective of cluster-dependent corralling by the cortical actin meshwork. Instead, our data suggest that clustering, or oligomerization more generally, increases the lifetime of membrane association, which in this system sets the timescales over which (1) the effects of advection are integrated and (2) flow-induced asymmetries are retained after flows cease. In this sense, oligomerization enhances a molecule’s effective memory of cortical flows in a manner consistent with recent theoretical analysis of the role of oligomerization in polarity circuits (Lang and Munro, 2022). Importantly, we have shown that modulation of this memory allows one to invert the polarity of a normally posterior PAR protein.

Whether this universal coupling of PAR proteins to cortical actin flows points towards a paradigm of bulk membrane flow remains unclear. None of the PAR proteins show obvious colocalization with actin and do not require actin to stabilize their localization to the membrane (Chang and Dickinson, 2022; Goehring et al., 2011a; Munro et al., 2004). However, lateral mobility of PAR-3 is enhanced by depolymerization of actin (Sailer et al., 2015) and it is possible that PAR proteins may nonetheless be molecularly linked to the actin cortex, perhaps through association into higher order, heterogenous assemblies with other proteins or lipids at the membrane:cortex interface. Resolving the nature of membrane flow and molecular features that enable advection in this system will require a more comprehensive survey of membrane-associated molecules.

Finally, how do our results impact our understanding of PAR-3 segregation? Through the use of a nanobody-based clustering strategy, Chang and Dickinson (2022) recently proposed an oligomer size threshold of three is required segregation. However, in our experiments, we found that dimers and tetramers were segregated with equal efficiency and even simply enhancing membrane association of monomers resulted in sufficient segregation to rescue division asymmetry, consistent with a primary role for membrane affinity in enabling segregation. However, one curious observation is that PAR-3 segregation was largely unchanged as a function of oligomer size for all oligomers > 1, despite size-dependent changes in membrane association as evidenced by membrane:cytoplasmic ratios. In other words, while increasing oligomerization enhanced the concentration of PAR-3 at the membrane it did not affect the relative fraction of membrane-associated PAR-3 that was segregated into the anterior. We suggest that oligomerization per se could enable PAR-3 to engage additional mechanisms that further stabilize asymmetry induced by cortical flows, thus explaining the lack of correlation between oligomer size and segregation efficiency. For example, in addition to stabilizing membrane association, oligomerization could also promote interactions with other molecules and/or enable some form of positive feedback (Dawes and Munro, 2011). While presumably not sufficient to drive polarization on its own, such feedback could reinforce advection-induced asymmetry to enhance and stabilize polarity (Lang and Munro, 2022). A key point of future work will be to determine how, if at all, such mechanisms combine with oligomerization-dependent membrane stabilization to enable high efficiency segregation of PAR-3.

## Conclusion

In conclusion, here we demonstrate that cortical flows can impact a broader range of membrane-associated molecules than previously anticipated. In the case of PAR polarity, this lack of selective advection requires that the membrane binding of PAR proteins be tuned to achieve efficient and selective transport during polarization.

## Methods

### C. elegans strains and culture conditions

*C. elegans* strains were maintained under standard conditions (Brenner, 1974) at 20°C (except where noted) on NGM (nematode growth media) plates seeded with OP50 bacteria. See Table S1 for details of all strains used in this work.

### Strain Construction

The insertion of HaloTag into the *par-1* locus was performed via CRISPR/Cas9 based on the published protocol (Dokshin et al., 2018). Briefly, tracrRNA (IDT DNA, 0.5 μl at 100 μM) and designed crRNA(s) for the target (IDT DNA, 2.75 μl at 100 μM) with duplex buffer (IDT DNA, 2.75ul) were annealed together (5mins, 95C) and then stored at room temperature until required. PCR products containing the DNA sequence to be inserted, i.e. HaloTag (sequence from (Dickinson et al., 2017), and HaloTag with 130 bp homology to the insertion site were generated and column purified (Qiagen, QIAquick PCR purification kit). PCR products were then mixed in equimolar amounts (2 μg each in a total of 10 μl) and annealed together (heated to 95°C and slowly cooled to room temperature) to generate a pool of products with long singlestranded DNA overhangs to act as the repair template. An injection mix containing Cas9 (IDT DNA, 0.5 μl at 10 mg/ml), annealed crRNA and tracrRNA along with the repair template was incubated at 37°C for 15 minutes before the debris in the mix was pelleted (15 mins, 14,500 rpm). Young gravid N2 adults were injected with the mix – mutants were screened by PCR and sequence verified. A *dpy-10* co-CRISPR strategy was used to facilitate screening (Arribere et al., 2014).

To generate point mutations/small insertions, mutation by CRISPR-Cas9 was performed based on the protocol published by (Arribere et al., 2014). To aid screening, silent mutations were engineered into the repair template to introduce a unique restriction site. Briefly, tracrRNA (IDT DNA, 0.5 μl at 100 μM) and crRNA(s) (IDT DNA, 2.7ul at 100uM) were combined in duplex buffer (IDT DNA, 2.8 ul), annealed together (5 mins, 95°C), and stored at room temperature until use. The final injection mix containing Cas9 (IDT DNA, 0.5 ul at 10mg/ml), annealed crRNA and tracrRNA along with the repair template for the mutation and a co-CRISPR marker (unc-58 or dpy-10) was incubated at 37°C for 15 minutes before the debris in the mix was pelleted (15 mins, 14,500rpm). Young gravid N2 adults were injected with the mix and mutants were selected, screened by PCR and/or restriction digest and sequence verified.

Primers and guides used to make all the strains generated in this project can be found in Table S1. Silent restriction sites can be identified in red in the table for the corresponding mutants.

### RNA interference mediated knockdown

RNAi was carried out using the feeding method (Kamath and Ahringer, 2003). RNAi NGM agar plates were prepared by seeding standard nematode growth media agar plates containing 1 mM IPTG (Isopropyl b-d-1-thiogalactopyranoside) with 150 μl of an overnight 3 ml culture of bacteria expressing the desired RNAi sequence (which was spiked with 30 μl of 1 M IPTG prior to seeding), and left at room temperature overnight. L4 larvae were placed on seeded RNAi plates for 16-36 hrs and incubated in a dark box at room temperature, unless stated otherwise.

### HaloTag Labeling

JF549-Halo and JF-646-Halo ligands were initially obtained from the laboratory of Dr. Luke Lavis (Grimm et al., 2015) and later purchased from Promega. Ligands were diluted in acetonitrile into 2 nmol and 0.5 nmol aliquots which were speedvac-ed (dry, 5 mins) to remove solvent, and stored at −20°C in a cardboard box. Labeling of the HaloTagged strains was broadly carried out as described (Dickinson et al., 2017) via liquid culture incubation. Briefly, overnight 2 mL OP50-LB cultures were centrifuged and the pellet was suspended in 200 μl of fresh S-medium solution (150 mM NaCl, 1 g/L K_2_HPO_4_, 6 g/L KH_2_PO_4_, 5 mg/L cholesterol, 10 mM potassium citrate pH 6.0, 3 mM CaCl_2_, 3 mM MgCl_2_, 65 mM EDTA, 25 mM FeSO_4_, 10 mM MnCl_2_, 10 mM ZnSO_4_,1 mM CuSO_4_). The Halo ligand was resuspended in 2 μl DMSO and added to the bacterial suspension to achieve a final concentration of 10 μM. 65 μl of this mixture was pipetted into 3 different wells in the center of a 96 flat bottomed well plate and 30-40 L4 worms were picked into each well. Water was added to surrounding wells to keep the plate from dehydration and the plate was covered in aluminum foil and placed in a 20° C shaking incubator overnight. Prior to imaging, worms were pipetted onto unseeded NGM plates. Once the liquid had settled, worms were picked onto new unseeded NGM plates and left for 15 minutes, and then onto OP50 or relevant RNAi plates.

### Live Imaging - Cortex and SPT

Embryos were dissected from gravid adult worms (picked as L4’s the night prior to the experiment) using a hypodermic needle into 7.5 μl of Shelton’s Growth Medium (SGM) containing 18.8 μm polystyrene beads (Polybead, Polyscience, Inc. Cat. # 18329). These were mounted between a glass slide and a high precision 1.5H 22 mm x 22 mm glass coverslip, with the edges sealed with wax (VALAP mixture, 1:1:1, Vaseline, lanolin, and paraffin wax). For worms incubated with Halo ligand, fresh glass coverslips were cleaned (20 min soaking in 96 % isopropanol with sonication, 3x rinsing with MilliQ H_2_O) and passivated with 30 μl 1 mg/mL PLL-PEG (SuSOS, Cat # PLL(20)-g[3.5]-PEG(2).20mg) for 40-60 minutes at room temperature. Prior to imaging, coverslips were rinsed in MilliQ H_2_O, dried on Whatman filter paper, and stored between lens cleaning tissue. Passivation is only effective for 48 hours.

Imaging was carried out with a 100x 1.49 NA TIRF objective on a Nikon TiE microscope equipped with an iLas2 TIRF unit (Roper), a custom-made field stop, 488 or 561 fiber coupled diode lasers (Obis), and an Evolve 512 Delta EMCCD camera (Photometrics), controlled by Metamorph software (Molecular Devices) and configured by Cairn Research. Filter sets were from Chroma: ZT488/561rpc, ZET488/561x, ZET488/561m, ET525/50m, ET630/75m, ET655LP. Images were captured in bright field, GFP/mNG (ex488/ZET488/561m), RFP/mKate/mCherry/Halo (ex561/ZET488/561m).

For advection/diffusion analysis, imaging was carried out at the onset of symmetry breaking. For single molecules (HaloTag), 2000 x 25 ms frames were streamed alternating between the GFP and RFP channels using a dual channel emission filter interleaved with 25 ms dummy channels, yielding a characteristic time interval of 100 ms. Bright field image sequences were captured before and after streams to ensure accurate staging and to monitor embryo viability. For clusters, a multi-dimensional acquisition was used to sequentially acquire images in the GFP, RFP and brightfield channels with 100 ms exposures and a 1 s time interval.

### Live Imaging - Midplane

Embryos used for midplane imaging were dissected 8 μl of egg buffer (118 mM NaCl, 48 mM KCl, 2 mM CaCl2, 2 mM MgCl2, 25 mM HEPES pH 7.4) containing 20.8 μm polystyrene beads (Polybead, Polyscience, Inc. Cat. # 18329) between a glass slide and a 1.5 22 mm x 22 mm glass coverslip and sealed with VALAP (1:1:1 vaseline, lanolin, petroleum jelly). Imaging was carried out with a custom X-Light V1 spinning disk system (CrestOptics, S.p.A.) with 50 μm slits, Nikon TiE with 63x or 100x objectives, 488, 561 fiber-coupled diode lasers (Obis) and Photometrics Evolve 512 Delta EMCCD camera, controlled by MetaMorph software (Molecular Devices) and configured by Cairn Research. Filter sets were from Chroma (Bellows Falls, VT): ZT488/561rpc, ZET405/488/561/640X, ET535/50m, ET630/75m. Multi-dimensional acquisition was used to sequentially acquire images in the following channels: DIC, GFP (ex488, ET535/50m), autofluorescence (ex488, ET630/75m), RFP (ex561, ET630/75m). To maximize signal to noise for segregation analysis, single images were captured at the end of cortical flow (defined as the departure of the male pronucleus from the posterior cortex) and after nuclear envelope breakdown, typically 5-6 minutes later.

### Image Analysis - Single Particle Detection and Tracking

Single molecule and cluster tracking were carried out in Python using the Trackpy package (https://github.com/soft-matter/trackpy). The custom Python code developed for the analysis is available at https://github.com/lhcgeneva/SPT. It implements the Crocker-Grier algorithm to localize particles to subpixel resolution in individual frames by fitting local intensity peaks to a Gaussian point spread function. Detection parameters such as the threshold intensity and diameter of the candidate particles are adjusted empirically for given imaging conditions. Particles are linked frame to frame, with additional user specified parameters. Parameters were optimized to minimize tracking errors and typically were as follows: feature size = 7 pixels, memory = 0 frames, minimum separation between features = 2 pixels, maximum distance features can move between frames = 4 pixels, minimum track length = 11 frames.

### Image Analysis - Particle Image Velocimetry

The local flow field of the acto-myosin cortex was measured by applying particle image velocimetry (PIV) to the NMY-2 image channel using the PIVlab MATLAB plugin (Thielicke and Stamhuis, 2014). Images were bleach corrected and rolling time averaged in Fiji (2 frames). The posterior region of interest was selected as an ROI, and images were pre-processed with a high-pass filter (size 10), with other preprocessing filters disabled. A FFT phase-space PIV algorithm with 2 passes (window sizes 64 and 32 pixels) was used with linear window deformation and a 2×3 Gaussian point fit. Postprocessing was done with two filters: (1) a velocity filter where manual velocity limits of 0.3 pixels/frame were drawn to remove any outliers and (2) a standard deviation filter where vectors that more than 5 standard deviations from the mean in each image were removed. Vectors that were removed were replaced by interpolation. On average 9.4% of vectors were replaced by interpolation (n = 31 embryos). The resulting flow fields have a vector every 16 pixels with additional intervening vectors generated by bicubic interpolation.

### Image Analysis - Segregation

Raw images were processed using SAIBR using N2 or NWG0038 animals for the autofluorescence calibration as required and ortical concentrations were obtained as described previously (Rodrigues et al., 2022). Briefly, a 50-pixel-wide (12.8 μm) line following the membrane around the embryo was extracted and computationally straightened, and a 20-pixel-wide (5.1 μm) rolling average filter was applied to the straightened image to reduce noise. Intensity profiles perpendicular to the membrane at each position were fit to the sum of a Gaussian component, representing membrane signal, and an error function component, representing cytoplasmic signal. Local membrane concentrations at each position were calculated as the amplitude of the Gaussian component and the anterior-most and posterior-most 30% of embryo perimeter averaged.

### Simulations - Single particle

Single particle stochastic simulations were carried out to simulate particle movement movies in 2D and benchmark algorithms relying on detection and tracking of bright fluorescent spots. A custom Python script was used for this purpose and is available at https://github.com/lhcgeneva/SPT. This script creates a series of images with diffusive particles overlayed on top of a noisy background. Spots are blurred by a Gaussian filter to simulate a microscope point spread function.

User defined variables include: the number of particles, diffusivity (in μm^2^/s), a coefficient of variation for diffusivity, time interval between subsequent frames (in seconds), total duration of ‘movie’ (in seconds), long axis & short axis to define area of each image (in μm).

The updated version of the simulation script for advection analysis also includes the additional user defined variables: dissociation rate (k_off_, in s^-1^) and advection velocities perpendicular to the long axis and parallel to the long axis (in μm s^-1^). Default values for each variable for a typical experiment have been defined in the script for easy future use.

When all variables have been defined, the script initializes by assigning particles a lifetime calculated from the probability distribution of the number of particles per frame for a given dissociation rate. The initial position of each particle is assigned as a set of random coordinates within the user defined area. A new position is created for each particle for each frame of its lifetime, by drawing from a probability distribution curve for steps in x and y directions, based on the particles defined diffusivity. If an advective component has been defined, the diffusive step is combined with its advective component. When the particle reaches the end of its lifetime, it is given a new random position and considered as a new particle appearing on the membrane. This association rate is equivalent to the defined dissociation rate and allows for the maintenance of a constant number of particles within any given frame. To prevent loss of particles due to boundary conditions, the final area of the images is determined by taking into account the maximum displacement of any particle and extending the image area on all sides by 6 pixels from the position of highest displacement.

### Simulations - Partial differential equations

To predict the efficiency of segregation, we modeled advection, diffusion and membrane exchange using a partial differential equation (PDE) model adapted from Goehring et al. (2011b). We used the following governing equations to model a single species (A):

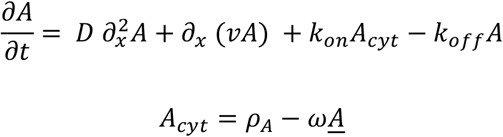

where A is the membrane concentration, A_cyt_ is the (uniform) cytoplasmic concentration, A^−^is the average membrane concentration, ψ is the surface area to volume ratio, ρ_A_ is the total amount of protein in the system, k_on_ and k_off_ are membrane binding and unbinding rates and D is the diffusion coefficient on the membrane. Systems were modeled as one-dimensional membranes 60 μm in length, where x is the position coordinate measured from the anterior pole. The cortical flow velocity (v) was specified to match an experimental flow profile as:

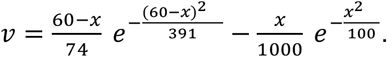

Values of k_off_ and D are specified in the main text. For all simulations we used ρ_A_ = 1.56 μm^-3^ and ψ = 0.174 μm^-1^ (Goehring et al., 2011b), and set kon equal to k_off_. Systems were initiated from a uniform equilibrium state and simulated using an adaptive Runge-Kutta scheme (Dormand and Prince, 1980) using custom code written in Python (available upon request - will be made available on GitHub).

### Quantification - Advection

Using SPT data, displacements were decomposed into local x and y axes defined as parallel and perpendicular to the local flow vector defined by PIV. For particles within a given bin (e.g. diffusivity or intensity), all individual displacements in x and y over a given time interval, τ, were pooled, plotted as a histogram and fit to a Gaussian distribution with advection (U_x_, U_y_) given by the mean displacements in x and y, respectively. Coupling coefficients were calculated as the ratio of U_x_ to the mean local velocity vector, v. For advection analysis, we let τ = 5 frames which provided a reasonable balance between maximizing the ratio of advection to diffusion, which increases with τ, and increasing noise due to the reduction in the number of displacements available, particularly for fast diffusing particles which tend to have shorter track lengths.

### Quantification - Diffusion

In general, mean particle diffusivity was calculated from particle step size, where <d^2^> is the mean square of all individual displacements for time delay t, with <d^2^> = 4D_ss_t for diffusion in two dimensions. For particles analyzed during the period of cortical flow, only the component perpendicular to the local flow field (Δy) was used, with <d^2^> = 2D_ss_t.

### Quantification - Segregation

Segregation was defined by several measures. Asymmetry index (ASI) is defined as (A - P) / (A + P), where A and P are the mean membrane concentrations in the anterior-most and posterior-most 30% of the embryo or simulation, respectively. Posterior depletion was defined as P/<I>, where P is as above and <I> is the mean membrane intensity. Decay times (t_1/2_) in simulations were fit to the values of ASI and Posterior depletion from the end of the cortical flow period.

## Contributions

Conceptualization: R.I., N.W.G.; Methodology: R.I., N.H., J.M., T.B., L.H.; Formal analysis: R.I., J.B.-P., J.M., T.B., K.N., N.W.G.; Investigation: R.I., N.H., J.B.-P., T.B., K.N., N.W.G.; Resources: R.I., J.B.-P., N.H., T.B.; Writing - original draft preparation: R.I., N.W.G.; Writing - review and editing: R.I., J.B.-P., N.W.G.; Supervision: R.G.E., N.W.G.; Project administration: R.G.E., N.W.G.; Funding acquisition: R.G.E., N.W.G.

## Acknowledgements

We thank Darius Koster and Guillaume Charras for comments on the manuscript, Mike Boxem for hosting R.I. to generate reagents, and Luke Lavis (Janelia Research Campus) for providing JF549-Halo and JF-646-Halo. Strains were graciously provided by Dan Dickinson and Ken Kemphues in advance of publication. Additional strains were provided by the Caenorhabditis Genome Center (CGC), which is funded by NIH Office of Research Infrastructure Programs (P40 OD010440). This work was supported by the Francis Crick Institute (N.W.G.), which receives its core funding from Cancer Research UK (FC001086), the UK Medical Research Council (FC001086), and the Wellcome Trust (FC001086), the EU Horizon 2020 Research and Innovation Program under the Marie Skodowska-Curie (grant no. 675407) (N.W.G.), and the Biotechnology and Biological Sciences Research Council (grant no. BB/M011178/1)(J.M., R.G.E).

This research was funded in whole, or in part, by the Wellcome Trust (FC001086). For the purpose of Open Access, the author has applied a CC BY public copyright license to any Author Accepted Manuscript version arising from this submission.

## Data Availability

Source code and documentation will be made available on GitHub.

## Supplemental Material

**Figure S1. Rescue of PAR-3(Δ69-82) by membrane targeting or oligomerization.**

**Figure S2. Segregation of PAR-6 under conditions of PAR-3-independent membrane loading.**

**Table S1. Strains and Reagents**

## Supplemental Movies

**Movie S1: Monomeric PAR-3 at the cortex undergoes advection PAR-3::Halo (CR1).** Near-TIRF timelapse image sequence of Halo::PAR-3(DeltaCR1). (Playback - 200X, Scale bar - 10 μm).

**Movie S2: Comparison of advection of Halo::PAR-3 variants.** Near-TIRF timelapse image sequence of Halo::PAR-3(WT) compared to RitC, 2mer and 4mer varants. (Playback - 200X, Scale bar - 10 μm).

**Movie S3: Advection is a general property of PAR proteins.** Near-TIRF timelapse image sequence of Halo fusions to PAR-1, PAR-2 and PAR-6. (Playback - 200X, Scale bar - 10 μm).

---

**Figure S1.**
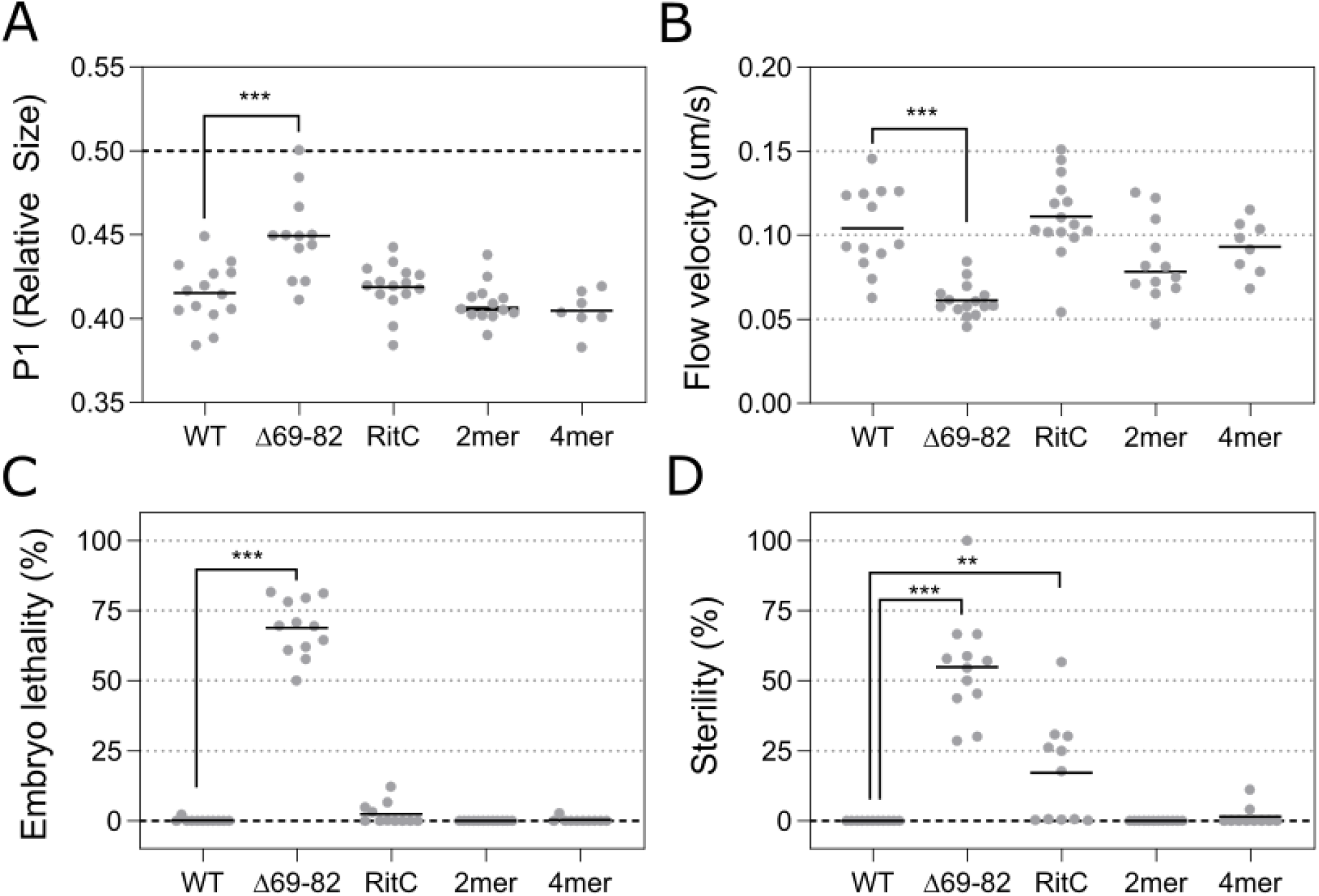
Rescue of PAR-3(Δ69-82) phenotypes by RitC, 2mer, and 4mer domains. **(A)** P0 division size asymmetry expressed as relative size of P1 as calculated from area of P1 and AB in a midsection image taken after cytokinesis (Area^P1^ / Area^P1+AB^). **(B)** Cortical flow velocity as measured by NMY-2 (PIV analysis). **(C-D)** Embryonic lethality (C) and sterility (D) of F1 progeny of animals homozygous for the indicated *par-3* allele. Significance vs. WT analyzed by one-way Anova with multiple comparison correction, with only p<0.05 shown. ***p<0.001, **p<0.01.

**Figure S2.**
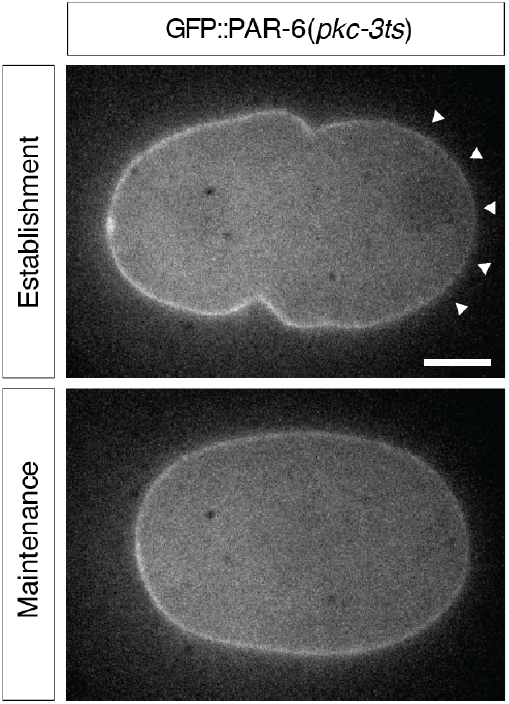
PAR-6 undergoes weak and transient asymmetry when decoupled from PAR-3 clusters. Representative midplane confocal images of PAR-6::GFP in wild-type or a pkc-3(ts) background in which PAR-6 membrane association is decoupled from PAR-3. When PAR-6 is not associated with PAR-3 clusters, segregation is weak and transient.

**Table S1.**
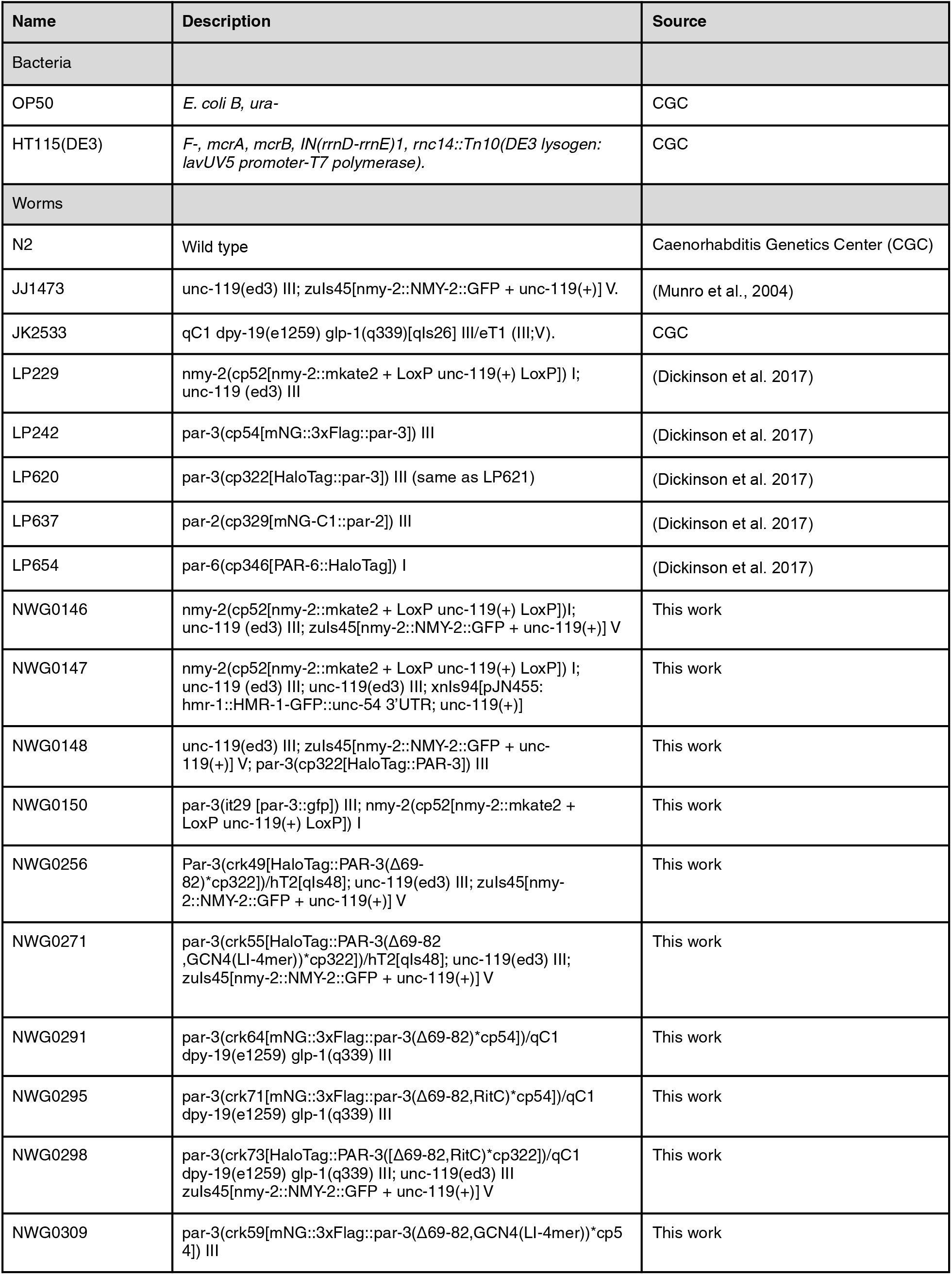
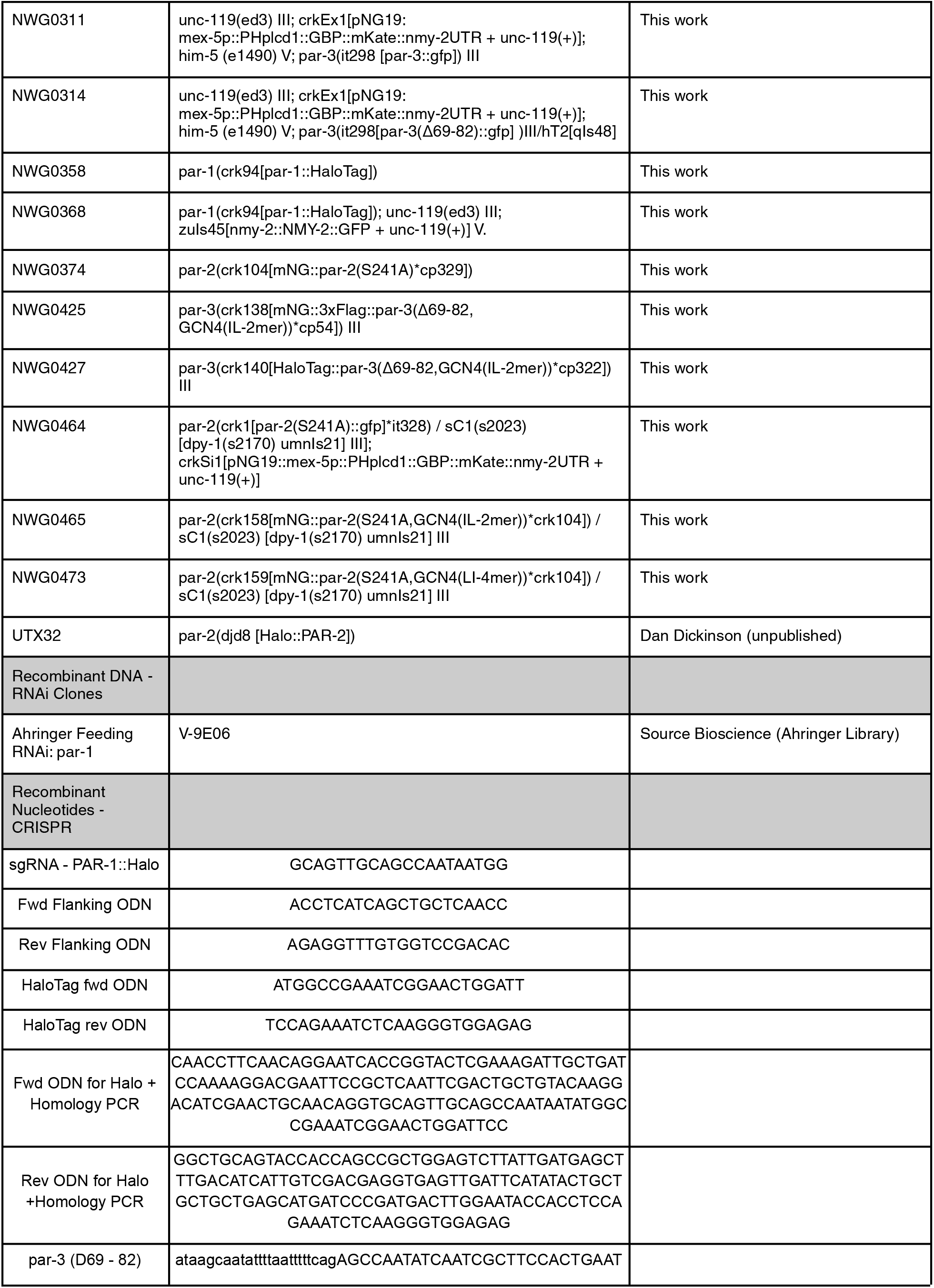
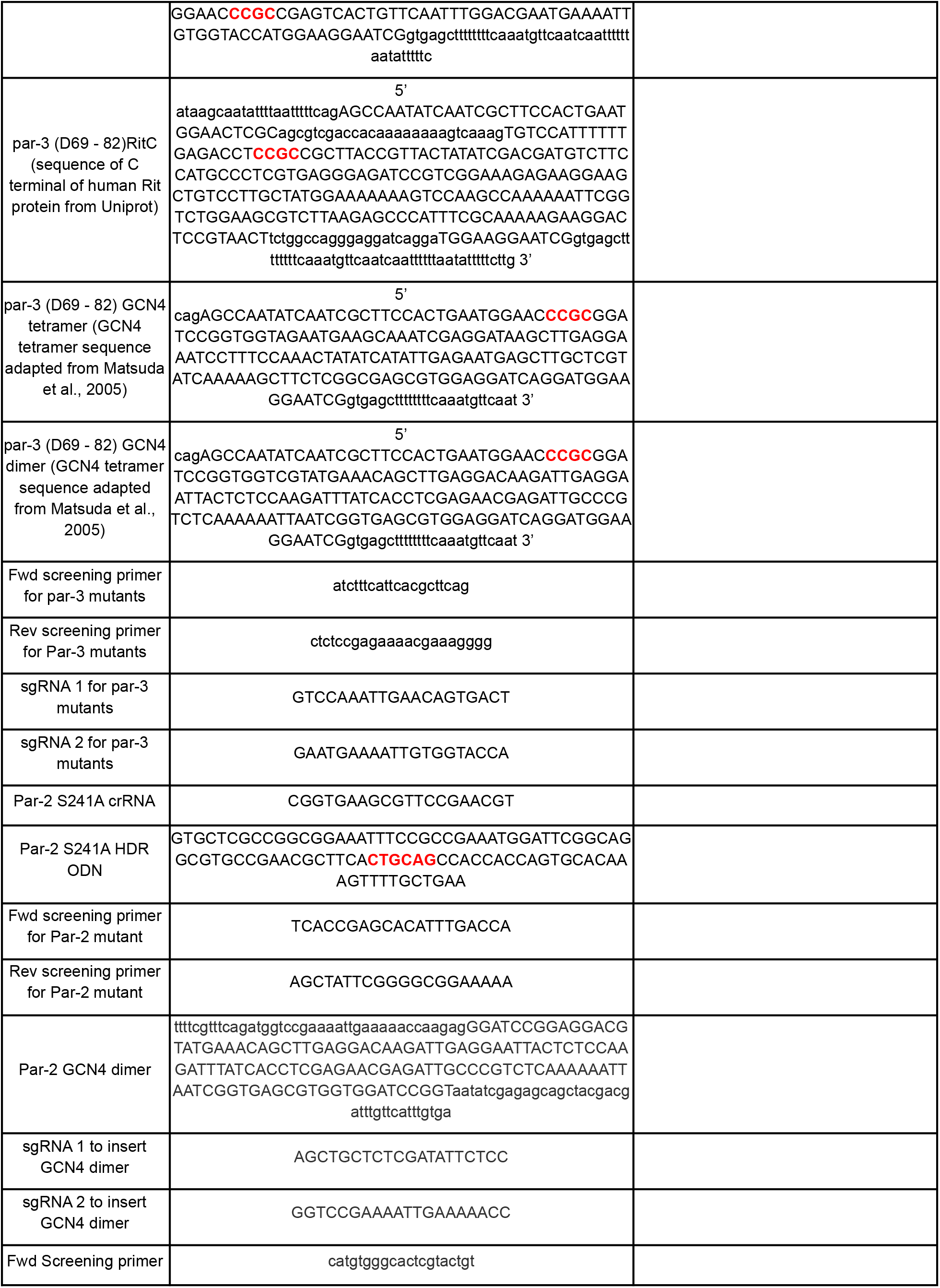
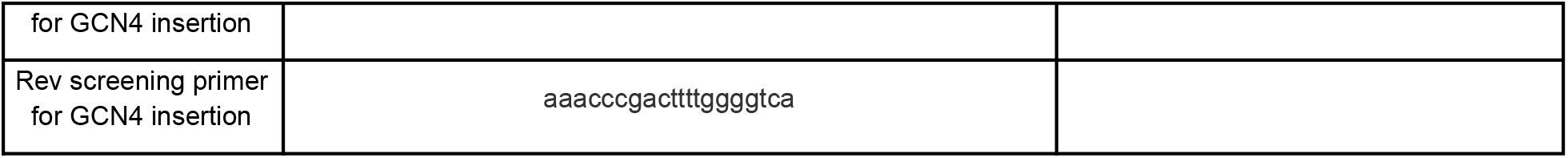
Strains and reagents.

